# Multi-platform discovery of haplotype-resolved structural variation in human genomes

**DOI:** 10.1101/193144

**Authors:** Mark J.P. Chaisson, Ashley D. Sanders, Xuefang Zhao, Ankit Malhotra, David Porubsky, Tobias Rausch, Eugene J. Gardner, Oscar Rodriguez, Li Guo, Ryan L. Collins, Xian Fan, Jia Wen, Robert E. Handsaker, Susan Fairley, Zev N. Kronenberg, Xiangmeng Kong, Fereydoun Hormozdiari, Dillon Lee, Aaron M. Wenger, Alex Hastie, Danny Antaki, Peter Audano, Harrison Brand, Stuart Cantsilieris, Han Cao, Eliza Cerveira, Chong Chen, Xintong Chen, Chen-Shan Chin, Zechen Chong, Nelson T. Chuang, Christine C. Lambert, Deanna M. Church, Laura Clarke, Andrew Farrell, Joey Flores, Timur Galeev, David Gorkin, Madhusudan Gujral, Victor Guryev, William Haynes Heaton, Jonas Korlach, Sushant Kumar, Jee Young Kwon, Jong Eun Lee, Joyce Lee, Wan-Ping Lee, Sau Peng Lee, Shantao Li, Patrick Marks, Karine Viaud-Martinez, Sascha Meiers, Katherine M. Munson, Fabio Navarro, Bradley J. Nelson, Conor Nodzak, Amina Noor, Sofia Kyriazopoulou-Panagiotopoulou, Andy Pang, Yunjiang Qiu, Gabriel Rosanio, Mallory Ryan, Adrian Stütz, Diana C.J. Spierings, Alistair Ward, AnneMarie E. Welch, Ming Xiao, Wei Xu, Chengsheng Zhang, Qihui Zhu, Xiangqun Zheng-Bradley, Ernesto Lowy, Sergei Yakneen, Steven McCarroll, Goo Jun, Li Ding, Chong Lek Koh, Bing Ren, Paul Flicek, Ken Chen, Mark B. Gerstein, Pui-Yan Kwok, Peter M. Lansdorp, Gabor Marth, Jonathan Sebat, Xinghua Shi, Ali Bashir, Kai Ye, Scott E. Devine, Michael Talkowski, Ryan E. Mills, Tobias Marschall, Jan O. Korbel, Evan E. Eichler, Charles Lee

**Affiliations:** Department of Genome Sciences, University of Washington School of Medicine, Seattle, WA, USA 98195; Molecular and Computational Biology, University of Southern California, Los Angeles, CA, USA 90089; European Molecular Biology Laboratory, Genome Biology Unit, Heidelberg, Germany; Department of Computational Medicine and Bioinformatics, University of Michigan, Ann Arbor, MI, 48109 USA; Center for Genomic Medicine, Massachusetts General Hospital, Department of Neurology, Harvard Medical School, Boston, MA, 02114 USA; The Jackson Laboratory for Genomic Medicine, Farmington, CT, 06032 USA; European Research Institute for the Biology of Ageing, University of Groningen, University Medical Centre Groningen, NL-9713 AV Groningen, The Netherlands; Center for Bioinformatics, Saarland University and the Max Planck Institute for Informatics, Saarbrücken, Germany; Institute for Genome Sciences, University of Maryland School of Medicine, Baltimore, MD, 21201 USA; Department of Genetics and Genomic Sciences, Icahn School of Medicine at Mount Sinai, New York, NY, 10029 USA; Xi’an Jiao Tong University, Shaanxi, China; Program in Bioinformatics and Integrative Genomics, Harvard Medical School, Boston, MA, 02115 USA; Department of Molecular Biophysics and Biochemistry, Yale University, 266 Whitney Avenue, New Haven, CT, 06520 USA; The University of Texas MD Anderson Cancer Center, Houston, TX, 77030 USA; Department of Bioinformatics and Genomics, College of Computing and Informatics, University of North Carolina, Charlotte, Charlotte, NC, 28223 USA; Department of Genetics, Harvard Medical School, Boston, MA, USA 02115 and the Stanley Center for Psychiatric Research, Broad Institute of MIT and Harvard, Cambridge, MA, 02142 USA; European Molecular Biology Laboratory, European Bioinformatics Institute, Wellcome Genome Campus, Hinxton, Cambridge, CB10 1SD, United Kingdom; Yale University Medical School, Computational Biology and Bioinformatics Program, New Haven, CT, 06520 USA; Biochemistry and Molecular Medicine, University of California Davis, Davis, CA, 95616 USA; UC Davis Genome Center, University of California, Davis, Davis, CA, 95616 USA; USTAR Center for Genetic Discovery & Department of Human Genetics, University of Utah School of Medicine, Salt Lake City, UT, 84112 USA; Pacific Biosciences, Menlo Park, CA, 94025 USA; Bionano Genomics, San Diego, CA, 92121 USA; Beyster Center for Genomics of Psychiatric Diseases, Department of Psychiatry University of California San Diego, La Jolla, CA, 92093 USA; Bioinformatics and Systems Biology Graduate Program, University of California, San Diego, La Jolla, CA, 92093 USA; 10X Genomics, Pleasanton, CA, 94566 USA; Illumina Clinical Services Laboratory, Illumina, Inc., 5200 Illumina Way, San Diego CA, 92122 USA; DNA Link, Seodaemun-gu, Seoul, South Korea; Department of Pediatrics, University of California San Diego, La Jolla, CA 92093 USA, Ludwig Institute for Cancer Research, La Jolla, CA, 92093 USA; High Impact Research, University of Malaya, 50603 Kuala Lumpur, Malaysia; TreeCode Sdn Bhd, Bandar Botanic, 41200 Klang, Malaysia; School of Biomedical Engineering, Drexel University, Philadelphia, PA, 19104 USA; Human Genetics Center, School of Public Health, The University of Texas Health Science Center at Houston, Houston, TX, 77225 USA; Department of Medicine, McDonnell Genome Institute, Siteman Cancer Center, Washington University School of Medicine, St. Louis, MI, 63108 USA; Department of Cellular and Molecular Medicine, University of California San Diego, La Jolla, CA, 92093 USA; Ludwig Institute for Cancer Research, 9500 Gilman Drive, La Jolla, CA, 92093 USA; Department of Computer Science, Yale University, New Haven, CT, 06511 USA; Institute for Human Genetics, University of California–San Francisco, San Francisco, CA, 94143 USA; Terry Fox Laboratory, BC Cancer Agency, Vancouver, BC, Canada V5Z 1L3; Department of Medical Genetics, University of British Columbia, Vancouver, BC, Canada V6T 1Z4; Molecular Neurogenetics Unit and Psychiatric and Neurodevelopmental Genetics Unit, Center for Human Genetic Research, Massachusetts General Hospital, Boston, MA, 02114 USA; Program in Medical and Population Genetics, Broad Institute of MIT and Harvard, Cambridge, MA, 02142 USA; Center for Mendelian Genomics, Broad Institute of MIT and Harvard, Cambridge, MA, USA 02142; Department of Human Genetics, University of Michigan, Ann Arbor, MI, 48109 USA; Howard Hughes Medical Institute, University of Washington, Seattle, WA, 98195 USA; Department of Graduate Studies – Life Sciences, Ewha Womans University, Ewhayeodae-gil, Seodaemun-gu, Seoul, South Korea 120-750

## Abstract

The incomplete identification of structural variants (SVs) from whole-genome sequencing data limits studies of human genetic diversity and disease association. Here, we apply a suite of long-read, short-read, and strand-specific sequencing technologies, optical mapping, and variant discovery algorithms to comprehensively analyze three human parent–child trios to define the full spectrum of human genetic variation in a haplotype-resolved manner. We identify 818,054 indel variants (<50 bp) and 27,622 SVs (≥50 bp) per human genome. We also discover 156 inversions per genome—most of which previously escaped detection. Fifty-eight of the inversions we discovered intersect with the critical regions of recurrent microdeletion and microduplication syndromes. Taken together, our SV callsets represent a sevenfold increase in SV detection compared to most standard high-throughput sequencing studies, including those from the 1000 Genomes Project. The method and the dataset serve as a gold standard for the scientific community and we make specific recommendations for maximizing structural variation sensitivity for future large-scale genome sequencing studies.

## INTRODUCTION

Structural variants (SVs) contribute greater diversity at the nucleotide level between two human genomes than any other form of genetic variation (Conrad et al. 2010; Kidd et al. 2010; Korbel et al. 2007; Sudmant et al. 2015). To date, such variation has been difficult to uniformly identify and characterize from the large number of human genomes that have been sequenced using short-read, high-throughput sequencing technologies. The methods to detect SVs in these datasets are dependent, in part, on indirect inferences (e.g., read-depth and discordant read-pair mapping). The limited number of SVs observed directly using split-read algorithms (Rausch et al. 2012; Kronenberg et al. 2015; Ye et al. 2009) is constrained by the short length of these sequencing reads. Moreover, while larger copy number variants (CNVs) could be identified using microarray and read-depth algorithms, smaller events (<5 kb) and balanced events, such as inversions, remain poorly ascertained (Sudmant et al. 2015; Chaisson et al. 2015).

One fundamental problem for SV detection from short-read sequencing is inherent to the predominant data type: paired-end sequences of relatively short fragments that are aligned to a consensus reference. The SV detection algorithms can thus be effective in unique sequences but break down within repetitive DNA, which is highly enriched for SVs (Conrad et al. 2010; Sharp et al. 2005). Another fundamental problem is that most SV discovery methods do not indicate which haplotype background a given SV resides on. Nevertheless, SVs are threefold more likely to associate with a genome-wide association study signal than single-nucleotide variants (SNVs), and larger SVs (>20 kb) are up to 50-fold more likely to affect the expression of a gene compared to an SNV (Chiang et al. 2017). Hence, SVs that remain cryptic to current sequencing algorithms likely represent an important source of disease-causing variation in unsolved Mendelian disorders and a component of the missing heritability in complex disorders (Manolio et al. 2009).

In this study, we sought to comprehensively determine the complete spectrum of human genetic variation in three family trios. To overcome the barriers to SV detection from conventional algorithms, we integrated a suite of cutting-edge genomic technologies that, when used collectively, allow SVs to be assessed in a haplotype-aware manner in diploid genomes. In addition, we also identified the optimal combination of technologies and algorithms that would maximize sensitivity and specificity for SV detection for future genomic studies.

## RESULTS

The goal of this study was to comprehensively discover, sequence-resolve, and phase all non-single-nucleotide variation in a selected number of human genomes. We chose three parent–child trios (mother, father and child) for comprehensive SV discovery: a Han Chinese (CHS) trio (HG00513, HG00512, HG00514), a Puerto Rican (PUR) trio (HG00732, HG00731, HG00733) and a Yoruban (YRI) Nigerian trio (NA19238, NA19239, NA19240). The Han Chinese and Yoruban Nigerian families were representative of low and high genetic diversity genomes, respectively, while the Puerto Rican family was chosen to represent an example of population admixture. The parents of each trio had been previously sequenced as part of the 1000 Genomes Project Phase 3 (1000 Genomes Project Consortium et al. 2015) and the children from each trio have been selected for the development of new human reference genomes (Chaisson et al. 2015). As a result, extensive genomic resources, such as SNV and SV callsets, single-nucleotide polymorphism (SNP) microarray data, sequence data and fosmid/bacterial artificial chromosome (BAC) libraries, have been developed to establish these trios as “gold standards” for SV assessment. We focused primarily on the three children for SV discovery using parental material to assess transmission and confirm phase.

We developed a multi-scale mapping and sequencing strategy using a variety of technologies to detect sequence variation of different types and sizes. To maximize sensitivity, we sequenced each child’s genome to a combined coverage of 223-fold (physical coverage of 582-fold) (Supplementary Table 1), using various short- and long-read technologies (Table 1). We discovered SVs using Illumina (IL) short-read whole-genome sequencing (WGS), 3.5 kb and 7.5 kb jumping libraries, long-read sequencing using PacBio^®^ (PB) (Menlo Park, CA) and optical mapping with Bionano Genomics (BNG) (La Jolla, CA). We also applied a series of genomic technologies capable of obtaining long-range phasing and haplotype structure: 10X Chromium (CHRO) (Pleasanton, CA), Illumina synthetic long reads (IL-SLR a.k.a. Moleculo), Hi-C (Lieberman-Aiden et al. 2009), and single-cell/single-strand genome sequencing (Strand-seq) (Falconer et al. 2012) technologies (Table 1; Supplementary Table 1; Supplementary Methods).

### Chromosomal level phasing and assembly of genomes

Assembly-based SV discoveries are usually represented as a single haplotype, rather than differentiating the two haplotypes of a diploid cell. This leads to reduced sensitivity for SV detection (Huddleston et al. 2017). We therefore aimed to resolve both haplotypes for the three children in this study by partitioning reads by haplotype and thereby detecting SVs in a haploid-specific manner. We applied WhatsHap (Martin et al. 2016) to IL paired-end, IL-SLR, and PB reads; StrandPhaseR (Porubsky et al. 2017) to Strand-seq data and LongRanger (Zheng et al. 2016) to CHRO data and compared them to more traditional trio-based (Martin et al. 2016) and population-based (Loh et al. 2016) phasing methods. As expected, the observed phased block lengths (Figure 1a) and marker densities (Figure 1b) differed substantially among the platforms but the amount of phasing inconsistencies, as measured by switch error rates (Figure 1c), was found to be very low (from 0.029% for CHRO to 1.4% for Hi-C). Since no single technology alone achieved the density, accuracy, and chromosome-spanning haplotyping necessary to comprehensively identify and assemble SVs throughout the entire human genome (Edge et al. 2017; Ben-Elazar et al. 2016; Porubský et al. 2017), we systematically evaluated the performance of all possible combinations of technologies. When combining a dense, yet local, technology (such as PB or CHRO) with a chromosome-scale, yet sparse, technology (such as Hi-C or Strand-seq), we obtained dense and global haplotype blocks (Figures 1d,e; Supplementary Material). To verify the correctness of chromosome-spanning haplotypes, we computed the mismatch error rates between the largest block delivered by each combination of technologies and the trio-based phasing (Figure 1f). The combination of Strand-seq and CHRO data showed the lowest mismatch error rate (0.23%), while phasing 96.5% of all heterozygous SNVs as part of the largest, chromosome-spanning haplotype block (Supplementary Table 2). We note that the switch error and mismatch rates are not constant for a given technology and can be influenced by factors such as sequencing coverage, data processing, and choice of restriction enzyme in the case of Hi-C.

**Figure 1.**
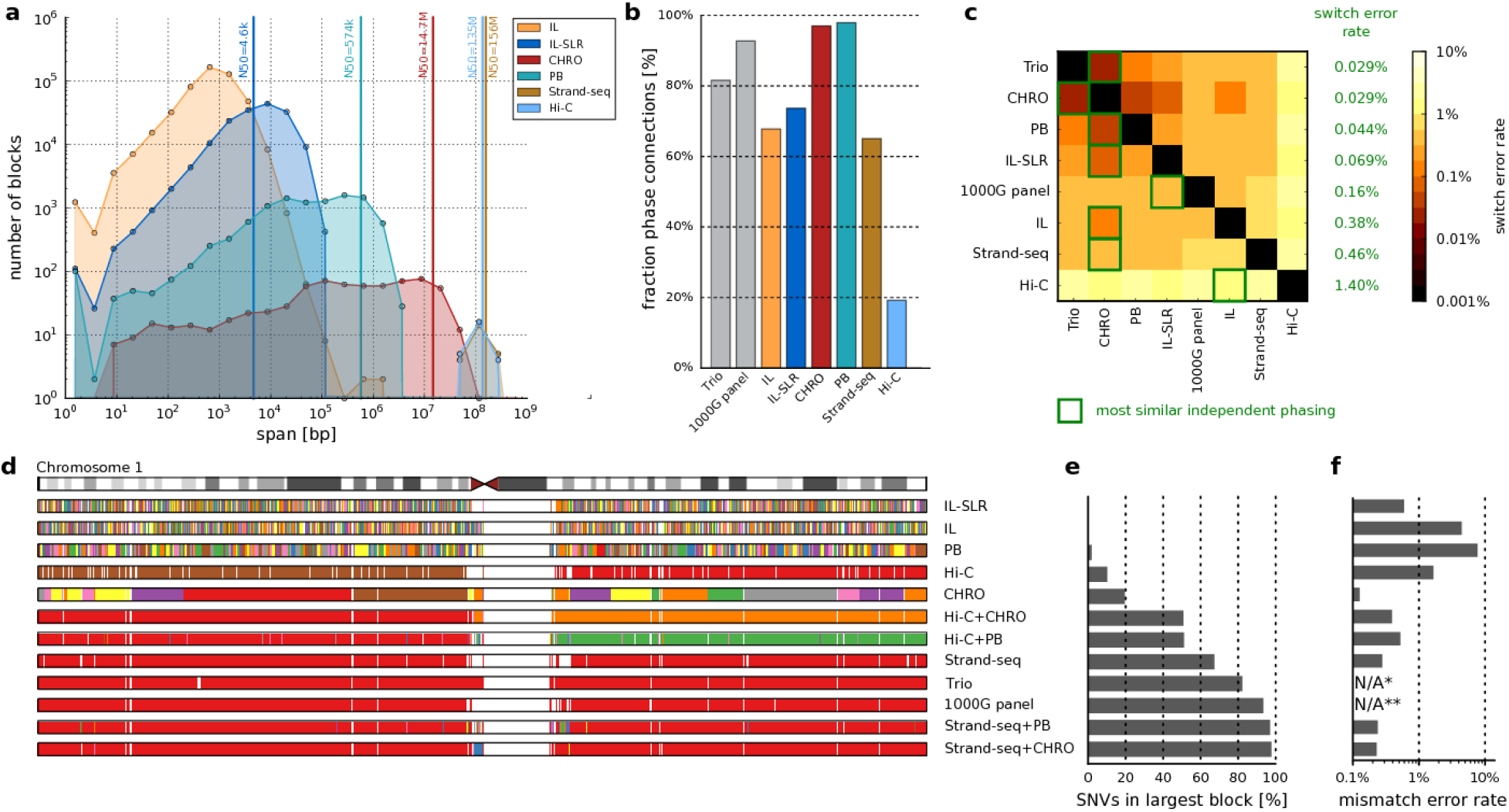
Characteristics of SNV-based haplotypes obtained from different data sources. (**a**) Distribution of phased block lengths for the YRI child NA19240. Note that Strand-seq haplotypes span whole chromosomes and therefore one block per chromosome is shown. For Illumina (IL) paired-end data, phased blocks cover <50% of the genome and hence the N50 cannot be computed. (**b**) Fraction of phase connection, i.e., pairs of consecutive heterozygous variants provided by each technology (averaged over all proband samples). (**c**) Pairwise comparisons of different phasings; colors encode switch error rates (averaged over all proband samples). For each row, a green box indicates the phasing of an independent technology with best agreement, with corresponding switch error rates given in green. (**d**) Each phased block is shown in a different color. The largest block is shown in red, i.e., all red regions belong to one block, even though interspaced by white areas (genomic regions where no variants are phased) or disconnected small blocks (different colors). (e) Fraction of heterozygous SNVs in the largest block shown in panel *d*. (**f**) Mismatch error rate of largest block compared to trio-based phasing, averaged over all chromosomes of all proband genomes (i.e., the empirical probability that any two heterozygous variants on a chromosome are phased correctly with respect to each other, in contrast to the switch error rate, which relays the probability that any two adjacent heterozygous variants are phased correctly). (*) Not available because trio phasing is used as reference for comparisons. (**) Not shown as population-based phasing does not output block boundaries; refer to Supplementary Material for an illustration of errors in population-based phasing.

Once chromosomal-level phasing was obtained for each child’s genome, we partitioned the PB reads according to haplotype. On average, 68% of reads could be haplotype-partitioned in each child (Supplementary Table 3). We then developed two complementary algorithms to assemble the haplotype-partitioned reads: an extension to the SMRT-SV method (Huddleston et al. 2016) (Phased-SV), which produced a separate assembly for each haplotype, and an extension to an assembly algorithm (Pendleton et al. 2015) (MsPAC), which combined separate haplotype-specific assemblies with *de novo* assemblies in autozygous regions (Supplementary Material). The assemblies covered, on average, 92.3% of the euchromatic genome (Supplementary Table 4) and produced contig N50 lengths ranging between 1.29 and 6.94 Mb (Supplementary Table 5). We then generated a high-quality consensus sequence for both assembled haplotypes (Chin et al. 2013) from which indels and SVs could be systematically discovered by mapping the contigs to the human reference.

In addition to providing a physical framework for phasing of all genetic variants, the parent–child trio data also allowed us to detect and refine meiotic breakpoints. Using Strand-seq data, meiotic breakpoints could be determined to a median resolution of less than 25 kb (Supplementary Material), which was further refined to ~1.5 kb (Supplementary Table 6) by the application of trio-aware phasing from PB reads (Garg et al., 2016). As expected, we observed an excess of maternal meiotic recombination events (Supplementary Material) (Broman et al. 1998; Hou et al. 2013; Kirkness et al. 2013; Lu et al. 2012). Further analysis of fine-mapped meiotic breakpoints support previously published results (Myers et al. 2008) of significant enrichment for L2 elements (p = 0.003). Similar enrichment was found for Alu retrotransposons (p = 0.003), especially the AluS class (p = 0.001) given the mere presence of these elements within the breakpoints (Supplementary Material). In addition, we identified an enrichment of a 15-mer motif at the breakpoints similar to previous studies (Myers et al. 2008) (Supplementary Material).

### Indel discovery (1–49 bp)

We generated a multi-platform indel callset by merging the IL- and PB-based callsets. Indels were detected in the IL WGS reads using GATK (DePristo et al. 2011), FreeBayes (Garrison and Marth 2012) and Pindel yielding, on average, 698,907 indel variants per child (Supplementary Material). The Phased-SV assembly alignments were used to detect indels >1 bp from the PB data yielding 345,281 indels per genome. The IL- and PB-based indel callsets showed similar size-spectrum distributions (Figure 2a) and were merged to yield, on average, 818,054 indels per individual. The unified indel callset showed the predictable 2 bp periodicity (Figure 2a; Supplementary Table 7) owing to the hypermutability of dinucleotide short tandem repeats (STRs) (Mills et al. 2006). The PB reads alone miss substantial numbers of IL-based indel calls (Figure 2c) and also lack the ability to reliably detect 1 bp indels, which are not included in the unified indel callset. However, more PB indels were discovered for variants greater than 15 bp (12% and 23% additional for insertions and deletions, respectively) (Supplementary Table 7). We were able to confirm 89% (529/594) of the homozygous sites (45% of all sites) that overlap with ~7 Mb of BACs from the children sequenced and assembled using high-coverage (>400x) PB reads (data not shown).

**Figure 2.**
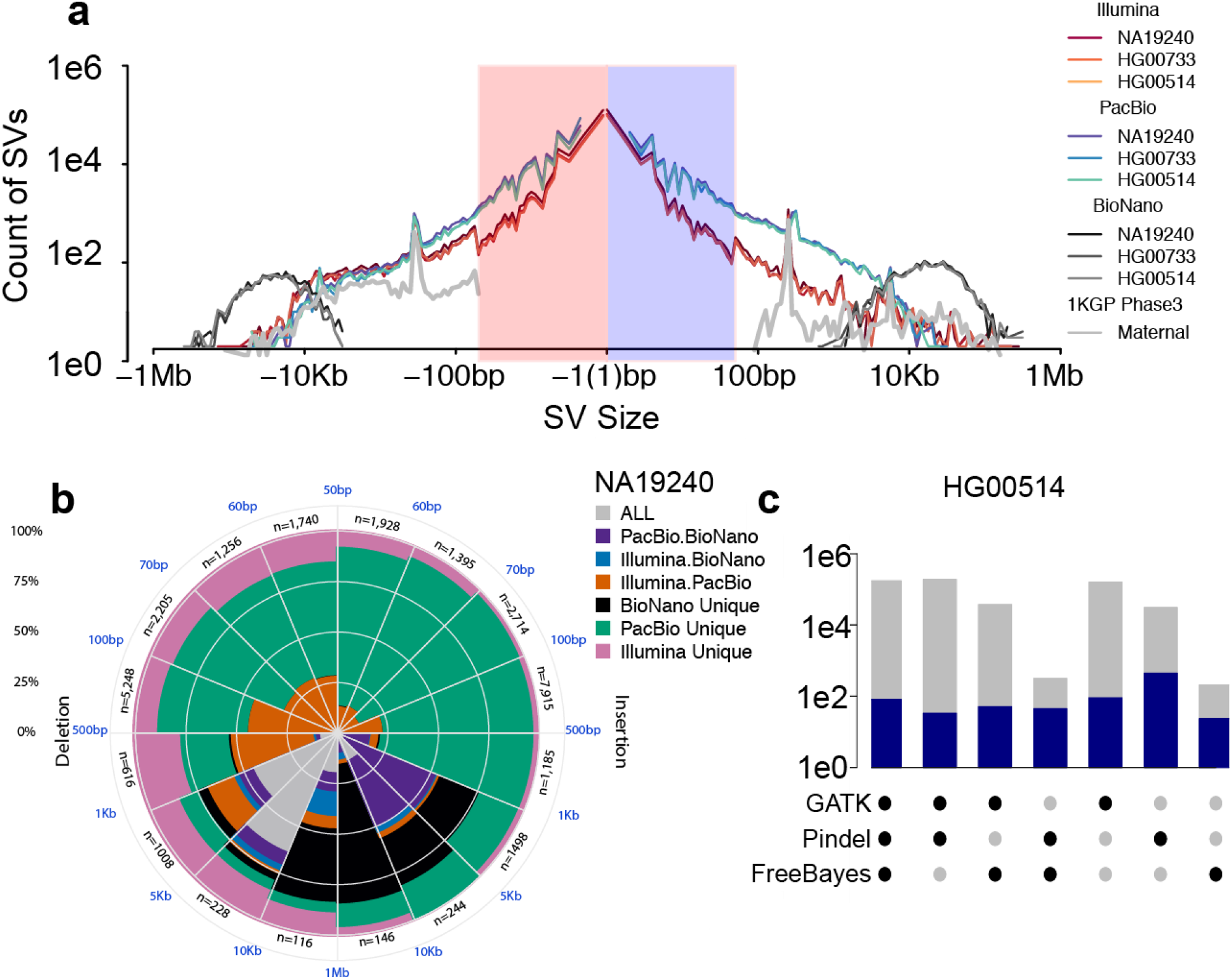
Comparison and integration of indel and SV callsets on HG00733, HG00514, and NA12940. (**a**) Length distribution of deletions and insertions identified by PB (blue), IL (red) and BNG (brown), respectively, together with averaged length distribution of SVs discovered in the maternal genomes by the 1000 Genomes Project Phase 3 report (silver). (**b**) Number of SVs discovered by one or multiple genome platforms in the YRI child NA19240. (**c**) Overlap of IL indel discovery algorithms, with total number of indels found by each combination of IL algorithms (gray) and those that overlapped with a PB-indel (blue) in the CHS child HG00514.

### SV discovery (≥50 bp)

We obtained a unified SV callset for each child from high-coverage IL WGS data, PB reads, and BNG assembly maps. To detect SVs in the IL data, we independently applied 13 SV detection algorithms (Supplementary Table 8), which included methods to capture paired-end, split-read, and read-depth information (Supplementary Material). Unlike the previous 1000 Genomes Project Phase 3 (1KGP-3) study, we sought to maximize discovery and did not strictly control for a given false discovery rate (FDR), opting to filter calls using orthogonal data in later steps. These calls were integrated into a unified Illumina-SV (IL-SV) callset (Supplementary Material) resulting in an average of 10,884 SVs per child (comprising of 6,965 deletions, 2,654 insertions and 814 duplications) and 20,395 nonredundant IL-SVs across all three children (Figure 2b). Approximately half (48.7%) of these IL-SV calls were annotated as high-confidence calls from a single algorithm, emphasizing the value of integrating multiple approaches from short-read sequencing data (Supplementary Material).

We generated a second set of SVs for each trio using the haplotype-resolved Phased-SV and MsPAC assemblies generated from the PB long-read sequencing data. Each assembly was mapped to GRCh38, and SVs were classified as insertions, deletions, and inversions. After applying a read-based consistency check (Supplementary Material) to remove assembly and alignment artifacts, the SVs from each assembly were merged into a per-individual unified callset (PB-SV). Excluding inversions, the integrated PB-SV callset consisted of an average of 24,825 PB-SVs per child (9,488 deletions and 15,337 insertions) for a total of 48,635 nonredundant PB-SVs across the three children (comprising 18,674 deletions and 29,961 insertions). The increase in sensitivity (threefold) from the PB-SV callset relative to the IL-SV callset was predominantly derived from improved detection of repeat-associated SV classes, particularly of intermediate-sized SVs (50 bp to 2 kb), and improved sequence resolution of insertions across the SV size spectrum. We note that the total SV count is dependent on the particular algorithm and gap penalties used because many of the SV calls (59% insertion, 43% deletion) map to tandem repeats where degenerate representative alignments are possible. For example, application of the double-affine gap penalty method NGM-LR (Sedlazeck et al. 2018) reduces the number of calls by 8.8% after similar call filtration and haplotype merging. More complex evolutionary models are necessary to determine the most biologically appropriate alignment parameters.

We validated the PB-SV callset genome-wide using three different approaches. First, we searched for evidence of each child’s SVs in the long-read sequencing data of one of the two parents (Supplementary Material). We determined that 93% of the homozygous SVs and 96% of the heterozygous SVs showed read support from one of the two parents consistent with SV transmission. Second, we genotyped the insertions and deletions against IL WGS data and found that, on average, of 91.7% (21,888) of Phased-SV calls could be genotyped (see below and Supplementary Material). Finally, we applied an orthogonal long-read technology, using Oxford Nanopore (ONT) to generate 19-fold sequencing coverage on the same PUR child genome with average read lengths of 12 kb. We also looked for support from individual ONT reads to validate the PB-SV calls. Requiring at least three ONT reads to support a PB-SV event, we achieved a 91% validation rate for SVs outside of tandem repeats and an 83% validation rate for SVs within tandem repeats (Supplementary Material).

A substantial fraction of human genetic variation occurs in regions of segmental duplication (Bailey and Eichler 2006), which are often missing from *de novo* assemblies (Chaisson et al. 2015). We compared the variation detected in regions of segmental duplication through read-depth to the segmental duplications resolved in the Phased-SV and MsPAC *de novo* assemblies. The haplotype-specific *de novo* assemblies overlapped 24.9% (43.6 Mb/175.4 Mb) of known human segmental duplications. The dCGH and Genome STRiP methods detect variation through changes in read-depth and are sensitive to copy number changes in highly duplicated regions. We determined that 93.8% and 73% of the copy number variable bases detected by dCGH and Genome STRiP, respectively, were not detected by *de novo* assembly (Supplementary Material). We also estimated that, on average, ~341 genes per child had at least one exon affected by a copy number change that was not detected in the *de novo* assemblies, highlighting the importance of continued read-depth-based CNV detection even when PB long-read-based *de novo* assemblies are generated.

### Characterization of inversions

Inversion variation has long been the focus of cytogenetic studies using karyotyping, and more recently, these large, rare inversions have been characterized at sequence resolution (Redin et al. 2017; Talkowski et al. 2012). However, submicroscopic and polymorphic inversions are ill defined by human genome sequencing, in part, because larger events tend to be flanked by virtually identical duplicated sequences that can exceed a million base pairs in length that cannot be bridged by short-read sequencing technology (Kidd et al. 2010). Moreover, the copy-neutral nature of simple inversions precludes detection by read-depth analysis. To generate a map of inversions across different length scales, we called inversions with five complementary techniques, including IL WGS, long-insert whole genome sequencing (liWGS), PB, BNG optical mapping, and Strand-seq. For Strand-seq, we developed a novel computational algorithm integrating inversion discovery with trio-aware phasing data to bolster accuracy (Methods) and only retained those calls that displayed haplotype support. A careful comparison of inversion calls revealed that Strand-seq was the only platform that made highly reliable calls on its own, while for the other technologies acceptable accuracy was achieved only for calls supported by at least two platforms (Figure 3a; Supplementary Material). The unified, high-confidence inversion callset comprised 308 inversions across the nine individuals, corresponding to an average of 156 inversions (~23 Mb) per genome. Of these 308, 75% were either primarily identified by Strand-seq (n = 170) or received additional Strand-seq genotype support (n = 59) (Methods). By comparison, 126 inversions in the unified callset were detected by IL WGS, 118 in PB, 91 in liWGS, and 28 in the BNG data.

**Figure 3.**
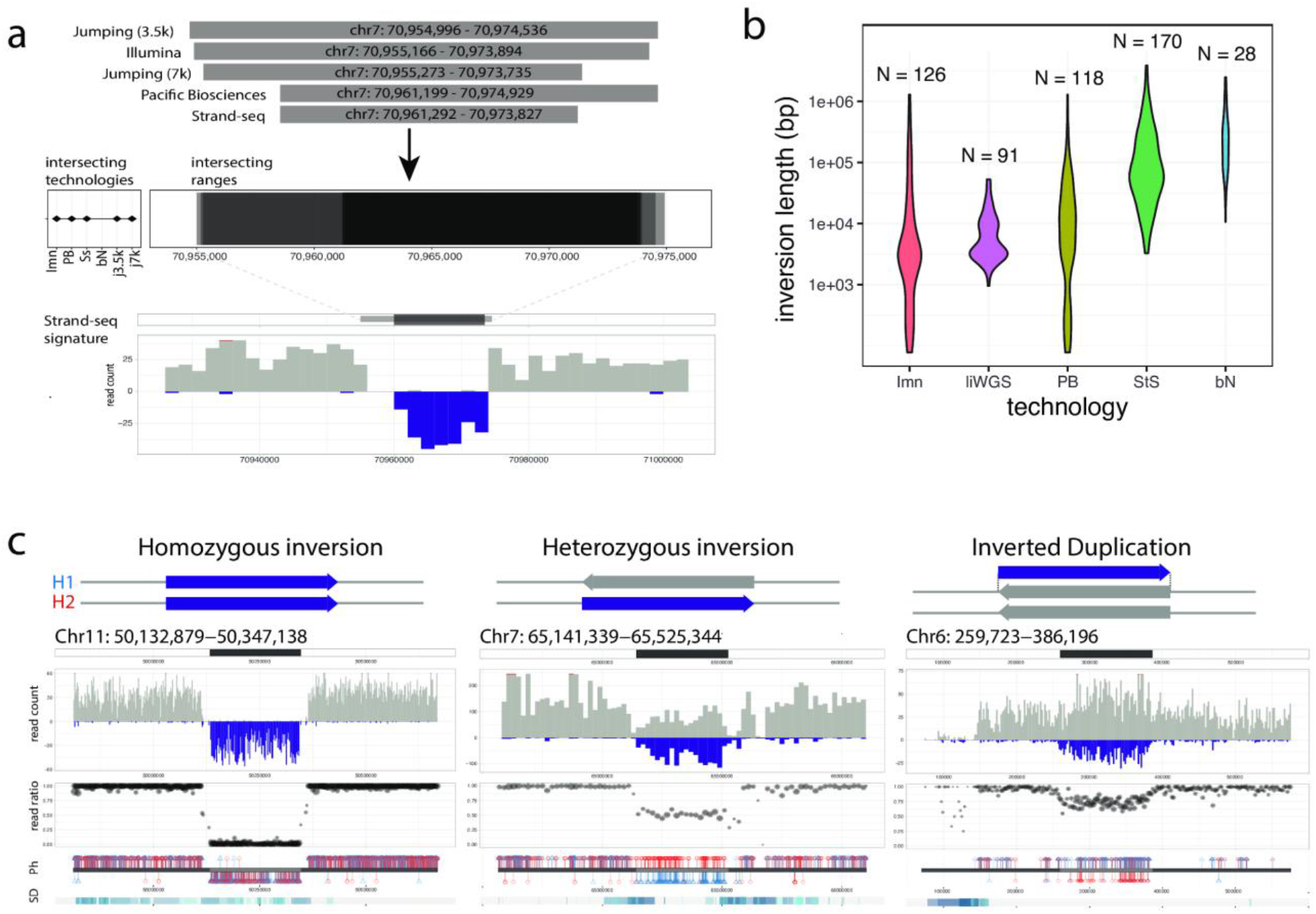
Characterization of simple and complex inversions. **(a)** Integration of inversions across platforms based on reciprocal overlap. Shown is an example of five orthogonal platforms intersecting at a homozygous inversion, with breakpoint ranges and supporting Strand-seq signature illustrated in bottom panels. **(b)** Size distribution of inversions included in the unified inversion list, subdivided by technology, with the total inversions (N) contributed by each listed. **(c)** Classification of Strand-seq inversions based on orthogonal phase support. Illustrative examples of simple (homozygous and heterozygous) and complex (inverted duplication) events are shown. Strand-seq inversions were identified based on read directionality (read count; reference reads in gray, inverted reads in purple), the relative ratio of reference to inverted reads within the locus (read ratio), and the haplotype structure of the inversion, with phased read data considered in terms of directionality (Ph; H1 alleles in red, H2 alleles in blue; alleles from reference reads are displayed above the ideogram and alleles from inverted reads are displayed below). Imn: Illumina. liWGS: long-insert whole genome sequencing libraries. PB: Pacific Biosciences. StS: Strand-seq. bN: Bionano. SD: segmental duplication. Ph: phase data.

The inversion size spectrum differed markedly among platforms (Figure 3b). IL WGS, PB, and liWGS excelled in mapping relatively small inversions (<50 kb), wherever breakpoint junctions could be traversed by DNA sequence reads. Indeed, the smallest inversions (<2 kb) were only detected by IL WGS and PB. In contrast, larger inversions (>50 kb) were nearly exclusively detected by Strand-seq. The Strand-seq technique offers the advantage of inversion detection solely by identifying DNA sequence strand switches internal to the inverted sequence, readily identifying inversions flanked by large segmental duplications that can be neither assembled nor traversed using standard DNA sequencing technologies (Sanders et al. 2016). Inversions called by Strand-seq show a median size of 70 kb (up to 3.9 Mb in length), in sharp contrast to IL-detected events, whose median size is 3 kb (down to 263 bp in length) (Supplementary Table 9).

Within the unified inversion callset, 73.7% (227/308) represent copy-neutral (i.e., simple) events, whereas 79 are more complex inversions containing embedded copy number variation (most in the form of inverted duplications). Consistent with previous observations that inversions map within segmental duplications, 50.7% of the inversions have both breakpoints mapping within segmental duplications (115/227)—an eight-fold increase when compared to unique regions of the genome. Furthermore, inversions within segmental duplications are ~20-fold larger, with a median length of 72.2 kb compared to a median length of 3.4 kb for inversions with breakpoints outside of segmental duplications. On average, each individual genome harbors 121 simple and 35 copy-variable inversions, approximately 2/3 (66.8%) are heterozygous and 1/3 (32.5%) are homozygous. Chromosomes 16 (5.2%), 7 (3.4%), X (3.3%) and 8 (3.0%) show the highest frequency of inversions, consistent with prior expectation (Sanders et al. 2016; Sudmant et al. 2015a; Chaisson et al. 2015). The inverted duplications typically exhibit highly variable copy number states, ranging between 0-10 (mean = 4) copies (Supplementary Table 10), indicating a large source of genetic variability between individuals. For instance, a 260 kb complex inversion mapping to chromosome 9 (at ~40.8–41.1 Mb) contains between 4-6 copies in each genome. Another notable example is an inverted duplication at the *DUSP22* locus (Figure 3c), for which a copy was known to be missing from the human reference (Genovese et al. 2013); we show it to be in the reverse orientation. Additionally, 40 inversions were found to be homozygous in all nine individuals and likely reflect minor alleles or remaining assembly errors in the human reference (Supplementary Table 11).

Inversion polymorphisms at several loci (such as 3q29, 7q11.23, 8p23, 15q13.3, 15q24, 17q12, and 17q21.31) were previously reported to predispose parental carriers to children with microdeletion and microduplication syndromes associated with developmental disorders (Koolen et al. 2008; Sharp et al. 2008; Antonacci et al. 2009). The substantially increased number of inversions generated in our study prompted us to investigate whether this association holds for other microdeletion and microduplication syndromes. Twenty-one events from our unified inversion callset displayed >80% overlap with one of the 255 “critical regions” (Weise et al. 2012; Coe et al. 2014) associated with genomic disorders (Supplementary Table 12). An additional 37 inversions were partially overlapping one of the 255 “critical regions”. Interestingly, in 12 cases we found novel inversions at both boundaries of the respective critical regions, including the 16p13.1 and 16p11.2-12 critical regions on Chromosome 16p (Extended Data Figure 1). A 1.9 Mb inversion at chromosome 3q29, for example, was previously shown to predispose to pathogenic SVs (Antonacci et al. 2009), and we identified a smaller 300 kb inversion intersecting the proximal breakpoint of this critical region. We hypothesize that the inversion polymorphisms we have identified will alter the orientation of low copy repeat sequences (see Extended Data Figure 1, right panel), and as such may differentially predispose individual loci to undergo pathogenic deletion or duplication via non-allelic homologous recombination.

### Mobile element insertions

Previous SV studies have been unable to resolve the sequences of large mobile elements in the human genome limiting our ability to assess differences in mutagenic potential between individual genomes. However, since PB long reads were routinely larger than 10 kb in length, we used the PB-SV callset to investigate not only the location but the sequence content of full-length L1 (FL-L1) elements. We detected an average of 190 FL-L1 elements with two intact open reading frames in the three children (Extended Data Figure 2; Supplementary Material). Only 56 of these copies are shared across the three genomes (Supplementary Table 13). This diversity in mobile element profiles likely influences L1 mutagenic potential. For example, while all three of the genomes are homozygous for one of the most active retrotransposon source L1 elements associated with human cancers (chr22:28663283, (Myers et al. 2008; Tubio et al. 2014)) and another L1 is highly active (i.e., “hot”) in the germline and cancers, each genome also harbors two to six unique hot L1 source elements. One of the unique hot L1 copies in the PUR individual is the *LRE3* element, which is the most active L1 source element in humans (Brouha et al. 2003, 2002). Twenty-eight FL-L1 copies with low-to-moderate levels of activity are also differentially present in the genomes of the three individuals. The cumulative differences in L1 mutagenesis that emerge from these diverse FL-L1 profiles suggest that, at a population level, such diversity may translate into differential risk levels for L1-mediated diseases such as cancers and other disorders (Tubio et al. 2014; Scott et al. 2016).

### Genotyping novel SVs in population cohorts

One of the advantages of having a more comprehensive set of sequence-resolved SVs is the ability to accurately genotype them in different human populations. We first genotyped SV calls from the base-pair-resolved PB-SV callset in a limited set of 27 high-coverage genomes using a sensitive but computationally intensive method, SMRT-SV Genotyper (Huddleston et al. 2017). An average of 91.7% (21,888) of Phased-SV calls could be genotyped with this approach across both insertions and deletions (Supplementary Material), with average Mendelian error rates of 14.1% for insertions and 8.7% for deletions. We also genotyped the small and large deletions from the PB callset with DELLY (Rausch et al. 2012) into a novel resource comprising 2,642 whole genomes derived from normal tissues sampled from cancer patients sequenced with IL reads (Waszak et al. 2017). With stringent filtering for high genotyping quality (≥35) and genotyping rate (≥75%), 13,233 total PB deletions could be genotyped across all samples. To assess the quality and effectiveness of our genotyping procedure, we made use of a statistically phased SNP set available for these 2,642 samples (Waszak et al. 2017) to assess the taggabillity of the genotyped deletions (Supplementary Material). We further evaluated the accuracy of our genotypes using principal component analysis revealing population structure, intensity rank-sum testing using SNP6 arrays available for 787 out of 2,642 samples (IRS FDR < 2%), and patterns of heterozygosity that confirmed an increased diversity in the samples of African ancestry (Supplementary Material). We next compared our PB-genotyped deletions to an IL-SV callset available for these 2,642 samples (Waszak et al. 2017) and to the 1000 Genomes Project phase 3 SV callset (Supplementary Material). This analysis revealed that a substantial fraction (34% and 43%, respectively) of the deletions at an SV size range of 25-500 bp are novel compared to these IL-SV callsets. In this size range, long-read derived *de novo* assemblies excel in sensitivity and a subset of these SV calls can be accurately genotyped using DELLY in IL data. While no SV showed extreme population stratification (FST > 0.85), there were 479 SVs with moderate differentiation (FST > 0.35), including nine deletions intersecting genes (Supplementary Table 14) and an exonic *CRIP2* deletion showing large population stratification (FST = 0.63; Supplementary Material). 113 deletions were in linkage disequilibrium (r^2^ > 0.8) to previously published genome-wide association study (GWAS) SNPs (MacArthur et al. 2017) (Supplementary Table 15) and 213 deletions tagged a blood eQTL SNP (r^2^ > 0.8) identified in the GTEx project (GTEx Consortium 2013), the latter of which included 14 deletions that were also in linkage disequilibrium with a GWAS hit. Notably, 54% (61 deletions) of the GWAS hits and 52% (110 deletions) in linkage disequilibrium to an eQTL were at the size range 25-500 bp where our study has improved sensitivity compared to IL-derived callsets (Supplementary Table 16). These new associations with variants of larger effect than SNPs, which represent only a fraction of the novel genetic variants discovered that could be genotyped, can now be pursued more systematically as potential pathogenic variants.

### Functional impact with respect to gene structure

An important consideration of increased sensitivity afforded by this multi-platform algorithm is its potential functional impact. Although the number of individuals compared is few, we sought to determine the number of genes that would be modified or disrupted based on the unified callset. On average, each child has 417 genes with indels and 186 genes with SVs predicted to affect protein-coding sequence. The coding frame is preserved for the majority of the genes with exonic indels (78.2%, 326/417) or SVs (30.1% (56/186) of intersecting genes (Supplementary Tables 17, 18). For frameshift events, we considered the intolerance to loss-of-function mutation as measured by pLI (Lek et al. 2016), which ranks genes from most tolerant (0) to least tolerant (1) of mutation. The median percentile of frameshift indels was 0.002 and SV was 0.155, indicating most of the loss-of-function variation is of modest effect (Supplementary Table 19). There were only two genes with high pLI (>0.90) that showed deletions concordant with the IL and PB data: *RABL1* (HG00514) and *ACTN1* (NA19240), and one 34 bp deletion leading to an alternate splice-site in *TSC2* in HG00733. We also identified 16 genes (such as *HTT*) that were intolerant to loss-of-function (pLI > 0.9) but carried in-frame indels—a potential source of triplet instability. 674 known canonical genes overlap inversions (Supplementary Table 20), of which 88% have isoforms entirely contained within the inversion (441), and 6% (32) are intronic. The remaining events are potentially gene disrupting, where three events overlap at least one exon of genes (*AQPEP, PTPRF*, and *TSPAN8*). Up to 55 genes have at least one isoform that spans one of the breakpoints of the inversion; however, the majority (95%) of these genes reside in segmental duplications where the exact breakpoint of the inversion cannot be easily resolved.

Variation in untranslated region (UTR) sequences can also affect gene expression leading to phenotypic consequences. We overlaid our SV dataset with UTRs and found that each child carried on average 155 genes with a deletion and 119 genes with an insertion in a UTR. Such genes, however, tended to be more intolerant to variation compared to exonic deletions, with a median pLI of 0.20. For example, there were 23 genes with UTR insertions or deletions with PLI scores >0.9: *ATP11A, BANP, BRWD3, DGKD, EIF4A3, FAM135B, FURIN, HCN1, IQSEC3, MEGF10, NIPBL, PAXBP1, PPP2R5E, SHB, SLC38A2, SNED1, SON, SREK1, TDRD5, TMEM165, XIAP, XPR1*, and *ZNF605*. The mean length of these UTR deletion variants was 176 bp and is similarly reflected in the technology bias for sensitivity; only one event was detected by BNG, nine by IL-SV, and 19 by PB-SV.

We also considered the overlap of SVs with functional noncoding DNA (fnDNA): specifically with 1.07M transcription factor binding sites (TFBS) and 2.86M conserved elements (CEs). Deletion variants overlapped an average of 861and 1,767 TFBS and 3,861 CEs in each child. However, small SVs rarely affected fnDNA: the average size of SVs that overlapped TFBS and CEs were 36,886 bp and 15,839bp, respectively. When considering the IL-SV and PB-SV callsets, the majority of TFBS and Ce deletions are detectable in the IL-SV callset (89.1% and 77.4%, respectively). Nevertheless, we estimate that 21 TFBS and 181 CEs would be missed per child by application of short-read sequencing technology alone. The opposite pattern exists for insertion SVs in fnDNA. While a smaller number of insertion SVs map inside TFBS (an average of nine per child, with average SV length of 665 bp) and 154 insertion SVs map inside a CE (average length of 930 bp), they were predominantly detected in the PB-SV callset; 57% of TFBS and 69% of CEs affected by SVs were detected only in the PB-SV callset compared to 7% and 5%, respectively, for IL-SV. Variants with imprecise insertion breakpoints, such as the BNG calls, were not considered. Thus, the application of multiple technologies enables additional resolution of smaller fnDNA SV.

### Platform comparisons and optimal indel and SV detection

The use of orthogonal technologies and various discovery algorithms on the same DNA samples provide an opportunity for a systematic assessment of the performance of individual as well as combinations of algorithms for indel and SV detection. While long-read technology generally outperforms IL-based algorithms for indel detection by ~50% for indels ≥15 bp, it is not reliable for single-base indels even at 40-fold sequence coverage, particularly in homopolymer regions. Benchmarking against the unified-indel dataset, we find that maximum sensitivity for IL indels requires application of three callers, including GATK, FreeBayes and Pindel (the latter of which has a higher false positive rate). Current large-scale studies rely mainly on IL sequencing, and computational resources limit the number of algorithms that can be applied to a genome. We therefore used a pan-SV callset (union of IL-SV, PB-SV, and BNG) to gauge the sensitivity and specificity of individual and combinations of IL-only algorithms. To construct the pan-SV callset, IL-SV insertions/deletions and BNG deletion calls were filtered according to orthogonal support datasets (formed from orthogonal callsets, raw PB reads, unfiltered PB-SV calls, and read-depth information). On average, 83% of IL, 93% of PB, and 95% of BNG deletion calls, and 82% of IL and 96% of PB insertion calls had orthogonal support. The concordant IL-SV and BNG calls were merged with the entire PB-SV callset to form the pan-SV callset. The unified callsets contained an average of 11,106 deletion and 16,386 insertion calls per individual (Table 2). As expected, the PB-SV callset provided the most unique calls, including 47% of deletions and 78% of insertions, which were primarily driven by tandem repeat variation (75%) and mobile element insertions (6.3%).

Across the entire IL-SV dataset, the deletion concordance to pan-SV was 82.9% (largely unaffected by size), whereas the insertion concordance to the pan-SV callset was 82.0%, decreasing in sensitivity with increased insertion SV length (Supplementary Table 21). The BNG mean concordance rate for deletions was 95.2%. When considering individual methods, the average concordance for deletion calls ranged from 46.4% to 99.3% with a median of 94.7% (Figure 4a), and for insertion calls ranged from <1% to 97% (Supplementary Material). When compared to the pan-SV callset, the concordant calls from individual algorithms detected 1.7%-40.7% of deletion and <1%-7.9% of insertion SVs.

**Figure 4.**
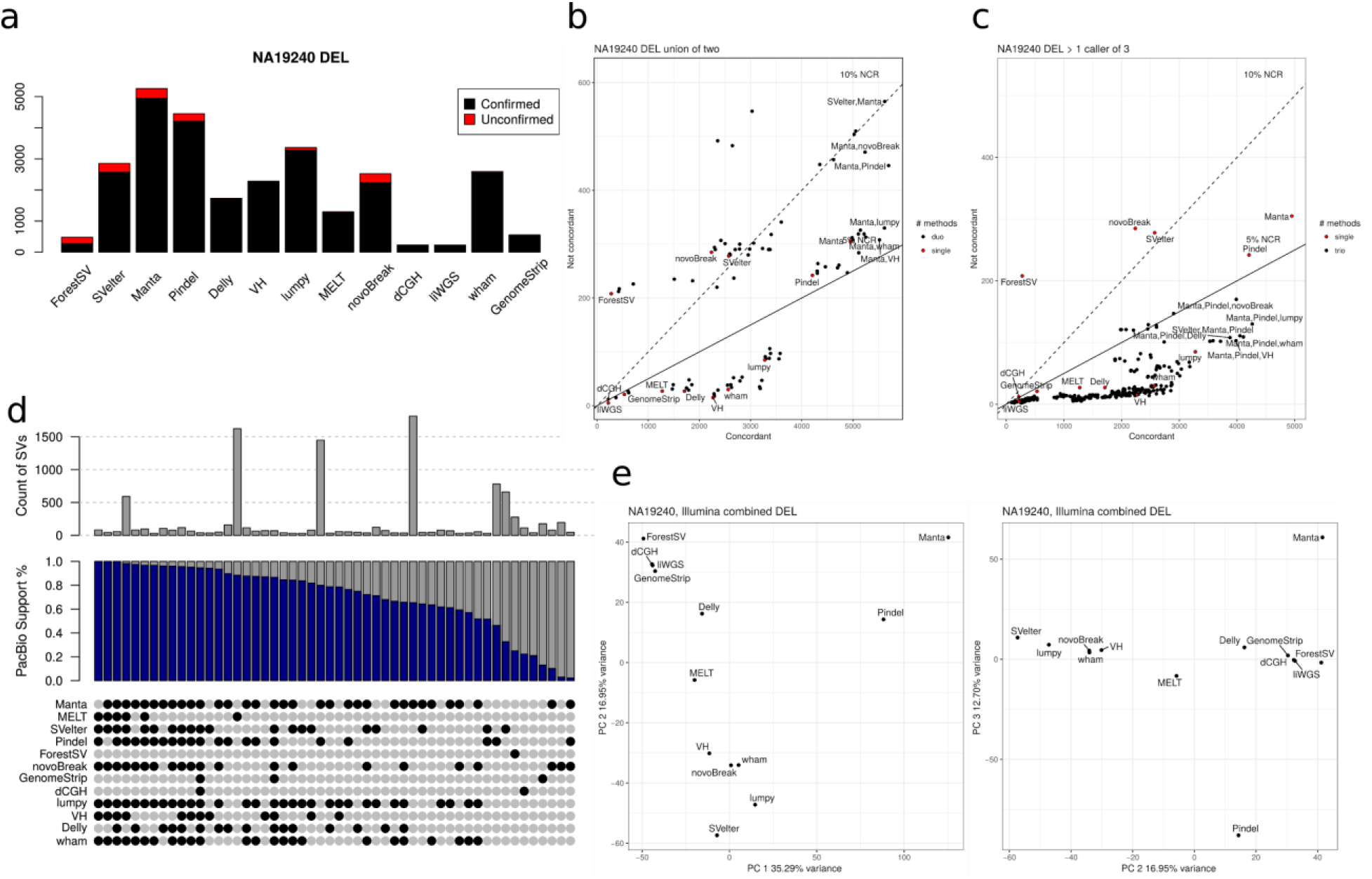
Concordance of IL methods compared against total IL callset and PB callset using orthogonal technologies. Results by algorithm shown for (**a**) the deletion concordance for individual methods, (**b**) the union of all pairs of methods, and (**c**) the requirement that more than one caller agree on any call. Individual callers are shown as red points for comparison. Pairs and triples of combinations are in black points. The solid and dashed lines represent the 5% and 10% non-concordance rates (NCR), respectively. The top five combinations of methods in each plot below the 10% NCR, along with the individual plots, are labeled. (**d**) Overlap of IL-SV discovery algorithms, with total number of SVs found by each combination of IL algorithms (gray) and those that overlapped with the PB-SV calls (blue) in the YRI child NA19240. (**e**) PCA of the genotypes of concordant calls of each method: PC 1 versus 2 (*left*), PC 2 versus 3 (*right*).

It has been shown previously that sensitivity for true SV calls (generated from IL datasets) can be improved by combining calls from more than one algorithm (Manolio et al. 2009; Mohiyuddin et al. 2015; Mills et al. 2011; Hehir-Kwa et al. 2016). Because it would be computationally burdensome for a large-scale WGS study to run all available algorithms, we used the integrated callset to compare how SV-calling performs under different combinations of algorithms (up to three) on standard IL data (i.e., without liWGS), testing both unions and intersections of callsets. We used the non-concordance rate (NCR) (1 - concordance) as a proxy for FDR. Considering the YRI child, NA19240, there were 29 combinations representing unions of two methods with an NCR less than 5% (Supplementary Material). For example, the union of the Pindel and VariationHunter callsets produced 4,892 calls at an NCR of 4.8%. Greater specificity may be obtained by requiring two of three methods to agree. There were 213 combinations of callsets with an NCR less than 2%, but the maximum sensitivity came from the combination of Manta, VariationHunter, and Lumpy (3,182 calls at an NCR of 2.9%) including 13 deletions from the pan-SV callset (including 65% of mobile element insertions and 4.5% of simple tandem repeats and variable-number tandem repeat deletions) (Figures 4b,c). The sensitivity of insertion-calling algorithms was driven by methods that detect mobile element insertions, particularly the MELT algorithm. We observed that while no single SV was called by every algorithm tested, there are often sets of algorithms that call similar variants (Figures 4d,e), and these PCA analyses may provide a conceptual framework for optimizing SV detection and computational burden in future studies.

## SUMMARY AND DISCUSSION

This study represents the most comprehensive assessment of SVs in human genomes to date. We employ multiple state-of-the-art sequencing technologies and methods to capture the full spectrum of genetic variation down to the single-nucleotide level, in a haplotype-aware manner. Our results indicate that with current methods, using multiple algorithms and datatypes maximizes SV discovery. The PB, Strand-seq, and CHRO data were combined to generate haplotype-resolved *de novo* assemblies constructed from phased PB reads. When paired with high-coverage IL sequencing and BNG SVs, we discovered approximately sevenfold more variation than current high-coverage IL-only WGS datasets (Sudmant et al. 2015), which is, on average, 818,054 indels (1-49 bp) and 27,622 SVs, including 156 inversions per person. Consistent with increased genetic diversity among African populations (1000 Genomes Project Consortium et al. 2015), we observed 20.7% more deletion and 9.4% more insertion variants in the Yoruban child than the Han Chinese child.

The long-read sequence data provided us with an unprecedented view of genetic variation in the human genome. Using average 5-10 kb reads at an average of 40-fold sequence coverage per child, we have now been able to span areas of the genome that were previously opaque and discover 2.48-fold more SVs than the maximum sensitivity achieved by integrating multiple algorithms for SV detection in short reads. Our analysis suggests that the majority (~83%) of insertions are being missed by routine short-read-calling algorithms. Specifically, the largest gain stems from tandem repeat and retro-transposon insertions in the 50 bp to 2 kb size range. Inversions represent another problematic class of human genetic variation. In 1KGP-P3, 786 inversions were identified across 2,504 genomes representing 3.3 Mb of sequence (Sudmant et al. 2015), and in the current study, we identified 308 inversions from just three family trios, totaling 36.4 Mb of sequence. This increase in sensitivity depended on the complementary nature of the five different technologies (Figure 3b). In the shorter size range, inversion discovery largely depended on a combination of IL and PB datasets, whereas for the larger events, Strand-seq was required. As a result, we were able to identify 181 inversions that were missed as part of 1KGP-P3. Most of these are large (>50 kb) constituting an average of 156 inversions and representing 22.9 Mb of inverted DNA per diploid genome, which corresponds to an ~480-fold increase in inverted bases per individual when compared to the 1000 Genomes Project (Sudmant et al. 2015). Our results indicate that for maximum sensitivity and specificity related to SV discovery it is essential to employ more than one detection algorithm and more than one orthogonal technology. This allowed us to locate new inversions to the boundaries of critical regions implicated in microdeletion and microduplication syndromes. Inversion polymorphisms have already been linked to disease, including Koolen-de Vries, Williams-Beuren, and 17q21 and 15q13.3 microdeletion syndromes (Koolen et al. 2008); (Sharp et al. 2008); (Antonacci et al. 2009). Here we nominate additional inversions that may similarly predispose critical regions to undergoing deletion or duplication.

It is not practical for large-scale studies to detect variation by employing the menagerie of sequencing methods and algorithms used in this study. Instead, these data serve as a guide for the trade-off between the cost of sequencing and desired sensitivity for SV detection. For example, we demonstrated entire chromosomal phasing using the Strand-seq and CHRO libraries; however, the Strand-seq method is not yet as widely implemented in sequencing facilities as Hi-C, which when combined with CHRO libraries provides chromosome-arm level phasing and is likely sufficient for many applications. Similarly, with the high-coverage IL sequencing and the many algorithms used here, it was possible to detect up to ~52% of the total number of deletion SVs and ~18% of insertion SVs. Moreover, we performed a series of down-sampling experiments to determine the equivalence of our analyses to datasets routinely used for large-scale studies (IL 30X coverage) and execution by non-expert users (i.e., by using default parameters) for SV detection (Supplementary Material). These analyses revealed just an 11% reduction in calls attributable to the lower 30X coverage, but a 23% reduction in calls using default parameters from the six algorithms with greatest contribution to our final callset. Collectively, these analyses suggest that if large-scale studies such as TOPMed (https://www.nhlbi.nih.gov) or CCDG (https://www.genome.gov) were to rely on an individual algorithm, we estimate the sensitivity to detect deletion SVs outside duplicated portions of the genome would be at most 40% with an FDR of ~7.6% (using Manta). Insertion sensitivity would fare far worse with an estimate of ~7% and an FDR of 4%, but only for mobile element insertions using MELT. While the majority of the variants associated with coding regions missed by IL-based analysis appear to be neutral in effect, there is a threefold increase of SVs detected in coding sequences for specific genes (albeit genes more tolerant to mutation) when including the PB-SV callset. Importantly, the addition of the PB-SV callset increases sensitivity for genetic variation, which could have a more subtle effect on gene expression changes, including a twofold increase in variation in UTR sequences and a ~20% increase of SVs detected in TFBS and CEs.

Our analyses indicate that the contribution of SVs to human disease has not been comprehensively quantified based upon studies that have relied upon short-read sequencing. Until the cost and throughput of long-read sequencing support larger-scale studies, we propose that future disease studies consider a triaged application of multiple technologies to comprehensively identify SVs. Families that have been sequenced using IL-based WGS should be analyzed using intersections of multiple SV-calling algorithms (e.g., Manta, Pindel, and Lumpy for deletion detection, and Manta and MELT for insertion detection) to gain a ~3% increase in sensitivity over individual methods while decreasing FDR from 7% to 3%. Because a disproportionate number of bases affected by variation occurs in segmental duplications and PB-based assembly does not resolve the segmental duplication regions entirely, there is a need to apply other algorithms, such as read-depth methods (e.g., dCGH or Genome STRiP), to detect changes in copy number in highly duplicated regions of the genome. The sequence structure of such variation is still not resolved and novel methods will need to be developed to sequence-resolve CNVs (M. J. Chaisson et al. 2017). Of note, there is a pressing need to reduce the FDR of SV calling to below the current standard of 5% because forward validation of all potentially disease-relevant events will be impractical at this threshold. We also predict that a move forward to full-spectrum SV detection using an integrated algorithm could improve diagnostic yields in genetic testing. Moreover, the proper application of SV detection for patient care requires a deeper understanding of germline SVs from more individuals across diverse global populations.

## DATA RELEASE

Underling sequencing read data from the various platforms can be accessed via the International Genome Sample Resource (IGSR) (Clarke et al. 2017) at http://www.internationalgenome.org/data-portal/data-collection/structural-variation. Indel variant calls will be made available with dbSNP build B151. SV calls are made available under dbVar accession nstd152.

## METHODS

See Supplementary Material.

## ACKNOWLEDGEMENTS

We thank Nancy Halsema and Karina Wakker-Hoekstra for help with preparing Strand-seq libraries, Lara Urban for assistance with eQTL characterization, and T. Brown for assistance in editing this manuscript. We also thank the people who generously contributed samples to the 1000 Genomes Project. Funding for this research project by the Human Genome Structural Variation Consortium (HGSVC) came from the following grants: National Institutes of Health (NIH) U41HG007497 (to C.L., E.E.E., J.O.K., M.A.B., M.G., S.A.M., R.E.M. and J.S.), NIH R01CA166661 (to S.E.D.), NIH R01HG002898 (to S.E.D.), NIH F31HG009223 (to E.J.G), NIH RO1HG008628 (to G.T.M.), NIH UO1HG006513 (to G.T.M.), NIH 1R21AI117407-01A1 (to A.B.), NIH R01HD081256 (to M.E.T.), NIH 1R01HG007068-01A1 (to R.E.M.), NIH RO1HG002385 (to E.E.E.), the R15HG009565 (to X.S.), the US Defense Advanced Research Projects Agency (N66001-15-C-4039 to X.S.), the Wellcome Trust grants WT085532 and WT104947/Z/14/Z and the European Molecular Biology Laboratory (to S.F., L.C., E.L., HZ.-B., P.F.), grant UM.0000125/KWJ.HI from the University of Malaya (to C.L.K.), by a National Health and Medical Research Council (NHMRC) CJ Martin Biomedical Fellowship (#1073726) to S.C., and an Advanced ERC grant (to P.M.L.). E.E.E. is an investigator of the Howard Hughes Medical Institute. J.O.K. is a European Research Council (ERC) investigator. C.L. is a distinguished Ewha Womans University Professor, supported in part by the Ewha Womans University Research grant of 2016-7.

## AUTHOR CONTRIBUTIONS

SV discovery: M.J.P.C., A.D.S., X.Z., A.M., D.P., T.R., E.J.G., O.R., L.G., Z.N.K., R.L.C., F.N., X.F., H.B., G.J. R.E.H., F.H., D.A., C.C., S.K-P., A.P., A.B., Z.C., K.Y.; PacBio assembly and analysis: M.J.P.C., O.R., A.M.W., C-S.C., A.B.; Strand-seq data generation and analysis: A.D.S., D.P., T.M., D.C.J.S., V.G., P.M.L., J.O.K.; Phasing: D.P., T.R., Y.Q., V.G., A.N., M.G., T.M.; Hi-C data generation and analysis: D.G., Y.Q., A.N.; Merging and analysis: M.J.P.C., A.D.S., X.Z., E.J.G., O.R., A.M., Z.N.K., R.L.C., X.F., P.A., S.C., A.M.W., A.H., X.C., D.A., M.G., B.J.N., S.M., C-S.C., P.M., X.Z-B., E.L., M.X., A.B., K.Y.; Genotyping: M.J.P.C., T.R., Z.N.K., P.A., S.Y.; Functional analysis: M.J.P.C., D.P., E.J.G., J.W., X.K., S.K., S.L., C.N., T.G., N.T.C., V.G., X.S.; Validation: A.D.S., R.E.H., F.H., S.C., D.L., A.F., J.Y.K., A.E.W., A.W.; Data production: A.D.S., S.C., A.S., K.M.M., C.C.L., C.L.K., Y.Q., E.C., W-P.L., M.R., C.Z., Q.Z., D.C.J.S., W.H.H., J.Y.K., J.E.L., J.F., J.L., S.P.L., K.V-M., G.R.; Data archiving: L.C., S.F., K.M.M., V.G.; Group organization: D.M.C., J.K., H.C., X.C., W.X., B.R., L.D., C.L.K., M.B.G., P.F., P-Y.K., P.M.L., G.M., W.X., K.C., J.S., X.S., A.B., K.Y., S.E.D., M.T., T.M., J.O.K., E.E.E., C.L.; Organization of supplementary material: M.J.P.C., X.Z., R.E.M.; Co-chairs: J.O.K., E.E.E., C.L.; Manuscript writing: M.J.P.C., A.D.S., E.J.G., A.M., J.S., S.E.D., M.T., T.M., R.E.M., J.O.K., E.E.E., C.L.; Display items: M.J.P.C., A.D.S., X.Z., E.E.G., T.M.

Phasing lead: T.M., Inversion lead: J.O.K. IL-SV co-leads: A.M., R.E.M.; PB-SV co-leads: M.J.P.C., A.B.; Indel lead: K.Y.; Hi-C lead: B.R.

---

**Extended Data Figure 1.**
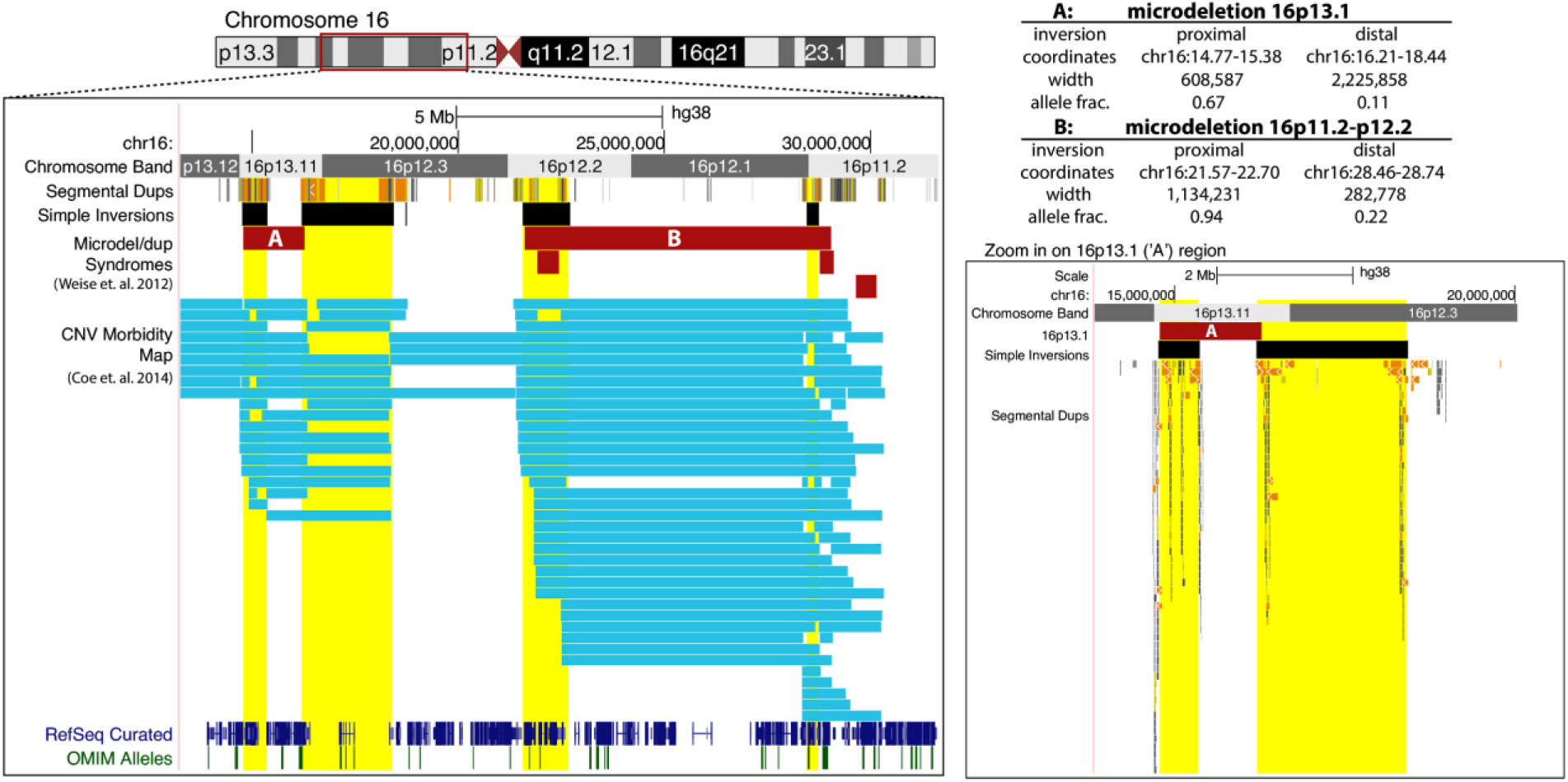
Examples of critical regions of microdeletion syndrome loci flanked by inversions. Genome browser view of two microdeletions located on chr16p (red bars, labeled ‘A’ and ‘B’) flanked by inversions located in the present study (black bars with yellow highlight). Expanded view of the segmental duplications rearranged by the inversions flanking region A are shown in the right panel. The coordinates, length, and genotypes for each inversion are listed in the table. Blue bars are microdeletions found in donors from the CNV morbidity map (Coe et al. 2014).

**Extended Data Figure 2.**
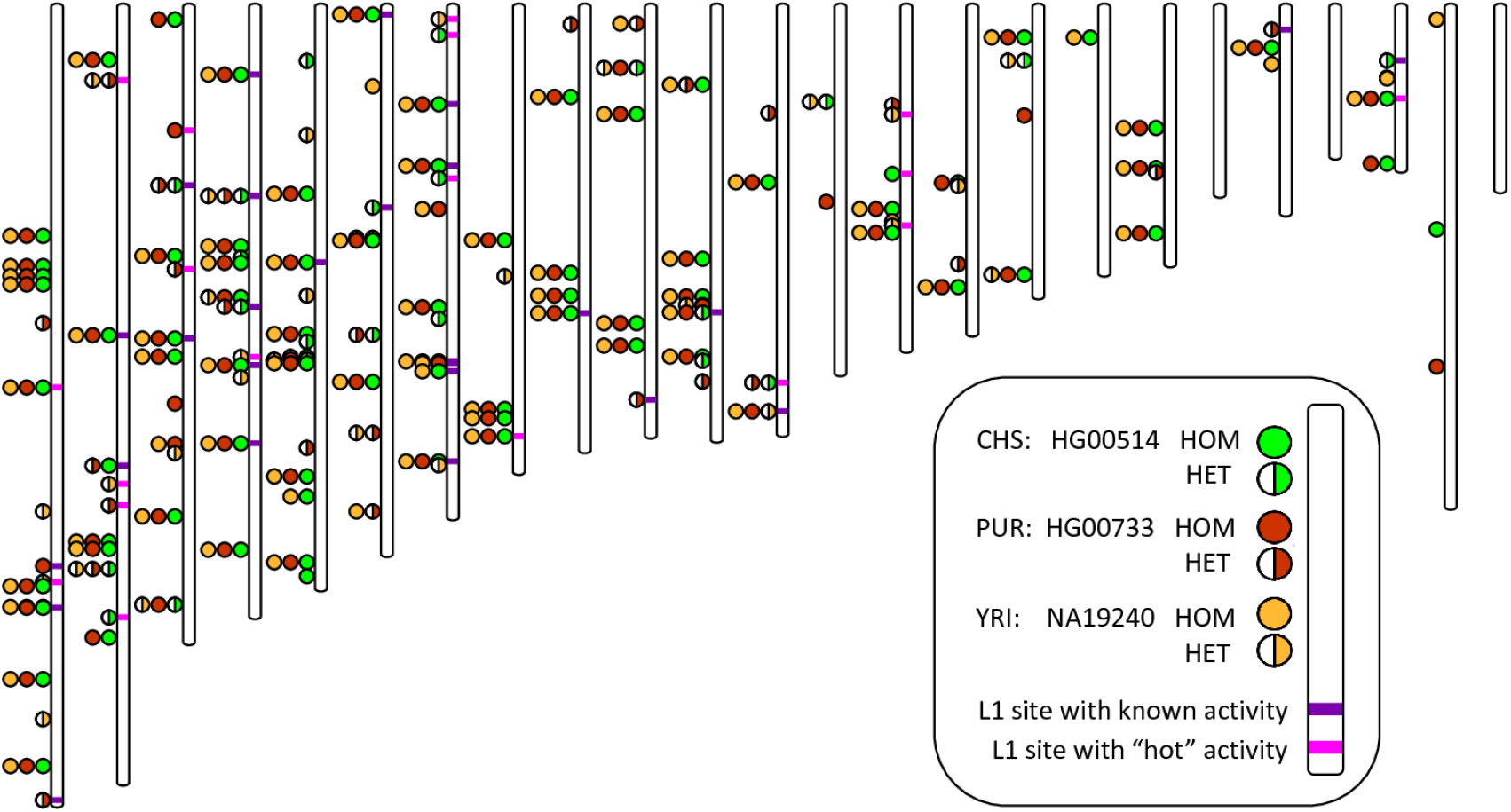
Intact FL-L1 source element profiles for the three children. Chromosome 1 through 22, X, and Y are displayed from left to right. FL-L1s with two intact open reading frames are represented by a circle in the color corresponding to the individual. The circle can either be filled or half-filled depending on the genotype in the respective individual. L1 sites with activity documented in the literature are depicted by light purple (highly active or “hot”) or dark purple (low to moderate activity) horizontal lines at the site.

